# Deletion of Robo4 worsens neuroinflammation and motor coordination in a mouse model of Alzheimer’s disease

**DOI:** 10.1101/2024.03.07.583582

**Authors:** Abigail E Cullen, Nick R Winder, Byron Lee, Sahana Krishna Kumaran, Nayantara Arora, Julia R Wolf, Grant D Henson, Randy L Woltjer, Ashley E Walker

## Abstract

Declines in vascular integrity are potential contributors to Alzheimer’s disease (AD) as these result in increased blood-brain barrier permeability and, as a consequence, accelerate neuroinflammation and cognitive impairment. Roundabout guidance receptor 4 (Robo4) is primarily expressed in endothelial cells and stabilizes the vasculature, and thus, has the potential to protect the brain in AD. To study the effect of Robo4 on neuroinflammation and cognitive function in the context of AD, we compared Robo4 knockout and wildtype mice crossed with mice with and without AD mutations (APP/tau). We found that the knockout of Robo4 led to greater astrocyte activation, as demonstrated by GFAP content, but this was dependent on the brain region studied. The knockout of Robo4 also led to greater activated microglia, as assessed by Iba1 content, but only in the presence of AD-related mutations. We found that AD mutations, but not Robo4, were associated with cognitive dysfunction measured by a nest-building test. In contrast, Robo4 deletion, but not AD mutations, was associated with impaired motor coordination. Lastly, Robo4 deletion was associated with greater arterial stiffness, but this trend did not reach statistical significance. In summary, these results demonstrate that Robo4 impacts neuroinflammation, motor coordination, and arterial stiffness, however, the impact on neuroinflammation is dependent on the presence/absence of AD-related mutations and the brain region examined.

## Introduction

It is becoming increasingly evident that vascular dysfunction is a contributor to Alzheimer’s disease (AD) and other dementias[1]. Featuring prominently in this vascular dysfunction is a decrease in vascular integrity [1]. In the brain, a decrease in vascular integrity leads to an increase in the permeability of the blood-brain barrier (BBB) that allows neurotoxic substances, inflammatory cells, and pathogens to enter the brain[1]. As such, greater BBB permeability is associated with increases in neuroinflammation, and neuroinflammation is further associated with the cognitive impairments and neuropathology that embody AD[2]. Indeed, greater BBB permeability is an early marker for cognitive decline[3]. However, the role of vascular integrity in AD is somewhat contradictory. While decreased integrity and increased BBB permeability have negative consequences, there is also evidence that signals that decreased integrity are associated with benefits. Central to these beneficial effects is vascular endothelial growth factor (VEGF), which increases permeability, but is also associated with positive outcomes in AD[4,5].Given the unclear role of vascular integrity in AD, a better understanding of these signaling pathways in the context of AD is needed.

The Roundabout family of transmembrane receptors has a role in vascular integrity. Roundabouts are best known for their role in axon guidance, and their primary ligands are Slit proteins[6].Roundabout guidance receptor 4 (Robo4) is primarily expressed in endothelial cells and stabilizes the vasculature[7]. Slit2, a ligand for Robo4, inhibits VEGF signaling in endothelial cells[7] and inhibits the vascular permeabilizing effects of pro-inflammatory cytokines, effects that are dependent on Robo4[8]. Slit2 is also known to inhibit endothelial cell migration and angiogenesis[8]. In summary, Robo4/Slit2 signaling increases vascular stability, and a decrease in this signaling increases vascular permeability.

Reductions in Robo4/Slit2 signaling are implicated in several brain-related impairments. For example, cerebral endothelial cells from old female mice have a lower Robo4 expression compared with their old male counterparts. It is suggested that this lower endothelial cell Robo4 expression increases the vulnerability to stressors, such as susceptibility to the cellular senescence caused by iron overload[9]. Additionally, brain endothelial cells from diabetic rats have lower Robo4 expression than cells from non-diabetic rats, and this decreased Robo4 expression is associated with detrimental excessive neovascularization[10]. Relatedly, Slit2 overexpression is associated with better cognitive performance, lower neuroinflammation, and better paravascular clearance in mice[11,12]. Administration of recombinant Slit2 also has benefits, demonstrated by lessening BBB permeability following surgical brain injury[13]. Thus, the Robo4 (and Slit2) pathway is known to have beneficial effects on the brain. However, the effects of Robo4 have not been studied in the context of AD. In particular, it is unknown if AD-related pathology (Aβ and tau) modulates the effects of Robo4.

To study the effects of Robo4 in the context of AD, we examined mice with Robo4 knockout (KO) crossed with mice with AD mutations (APPswe and tauP301L). We hypothesized that Robo4 KO mice would have greater neuroinflammation and worse cognitive function compared to Robo4 wildtype (WT) mice. We further hypothesized that there would be an interaction between Robo4 and AD mutations, such that the Robo4 KO x AD+ group would have worse neuroinflammation and cognitive function compared with both the Robo4 KO x AD- and Robo4 WT x AD+ groups.

## Results

### Robo4 deletion leads to astrocyte activation in AD mice

In the hippocampus, we observed no main or interaction effects of AD and *Robo4* genotype on GFAP expression (all p>0.05), indicating that in the hippocampus neither genotype influences the activation of astrocytes (**Figure 1A**).

**Figure 1.**
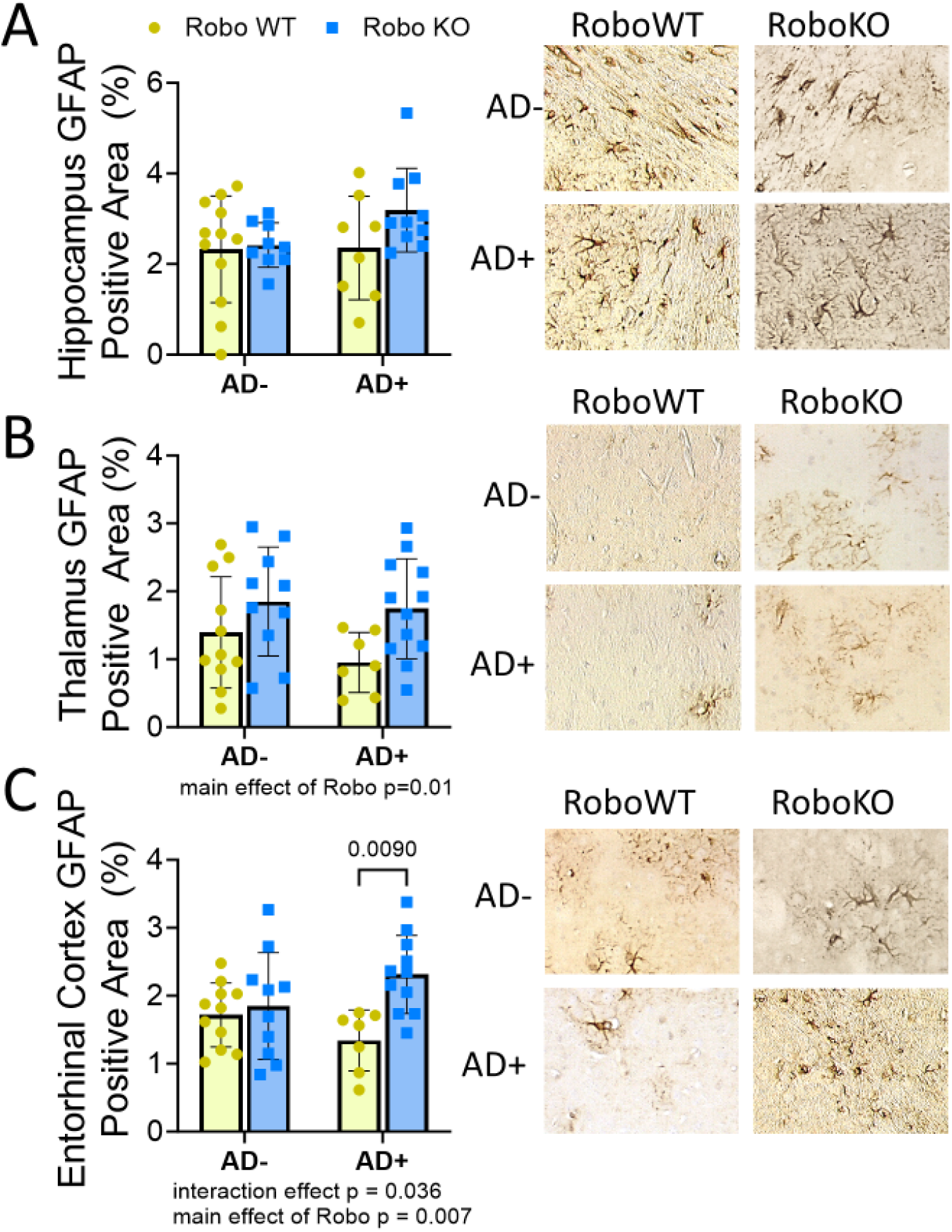
Astrocyte content is influenced by *Robo4* and AD genes. GFAP quantified by immunohistochemistry in the (A) hippocampus, (B) thalamus, and (C) entorhinal cortex for Robo4 wildtype (WT) or knockout (KO) mice that had mutant Alzheimer’s disease (AD) genes present (+) or absent (-). (n=7-12/group). Neither *Robo4* or AD genes caused significant differences in GFAP content in the hippocampus. Conversely, there was a main effect of *Robo4* genes on GFAP content in the thalamus and entorhinal cortex (p=0.01 and 0.007, respectively). There was an interaction effect of AD and *Robo4* in the entorhinal cortex (p=0.036). Representative images to the right. Values are mean±SD.

In the thalamus, there was a main effect of *Robo4* on GFAP (p=0.01) (**Figure 1B**). In contrast, we observed a main effect of AD on vimentin expression in the thalamus (p=0.02) (**Supplemental Figure 1B**). These results suggest that the effects of AD and *Robo4* on astrocytes in the thalamus depend on the marker measured.

In the entorhinal cortex, we observed similar trends for GFAP as in the thalamus (**Figure 1C**). There was a main effect of *Robo4* genotype on GFAP expression in the entorhinal cortex (p=0.007). In the AD mice, *Robo4* KO mice had more GFAP expression compared with the *Robo4* WT mice (p=0.009).

The data for vimentin expression in the hippocampus and entorhinal cortex were not normally distributed after transformation and were therefore not analyzed. The transformed data can be found in **Supplemental Figure 1A and C**.

### Robo4 deletion leads to microglia activation in AD mice

In the hippocampus and thalamus, there was an interaction effect between *Robo4* and AD genotype on Iba1 expression (p=0.01 and p<0.0001, respectively) (**Figure 2 and B**). There was a trend for lower Iba1 expression in the hippocampus of *Robo4* KO animals compared to their *Robo4* WT counterparts (p =0.06). There was significantly reduced Iba1 expression in the thalamus of *Robo4* WT x AD animals compared to *Robo4* WT (p=0.02). In *Robo4* KO, Iba1 expression was significantly reduced compared to their *Robo4* KO x AD counterparts (p=0.008). In the absence of AD genes, Iba1 expression was significantly lower in *Robo4* KO mice compared to *Robo4* WT (p=0.003). Conversely, in the presence of AD genes, *Robo4* KO mice showed significantly higher Iba1 expression compared to *Robo4* WT. In contrast to the thalamus, there were no additional differences detected in the hippocampus (all p>0.05). While there were interaction effects in the hippocampus and thalamus, there were no significant interactions or main effects for Iba1 in the entorhinal cortex (**Figure 2C**). These results indicate that *Robo4* has a differential impact on microglia activation dependent on the presence of mutant AD genes and location within the brain.

**Figure 2.**
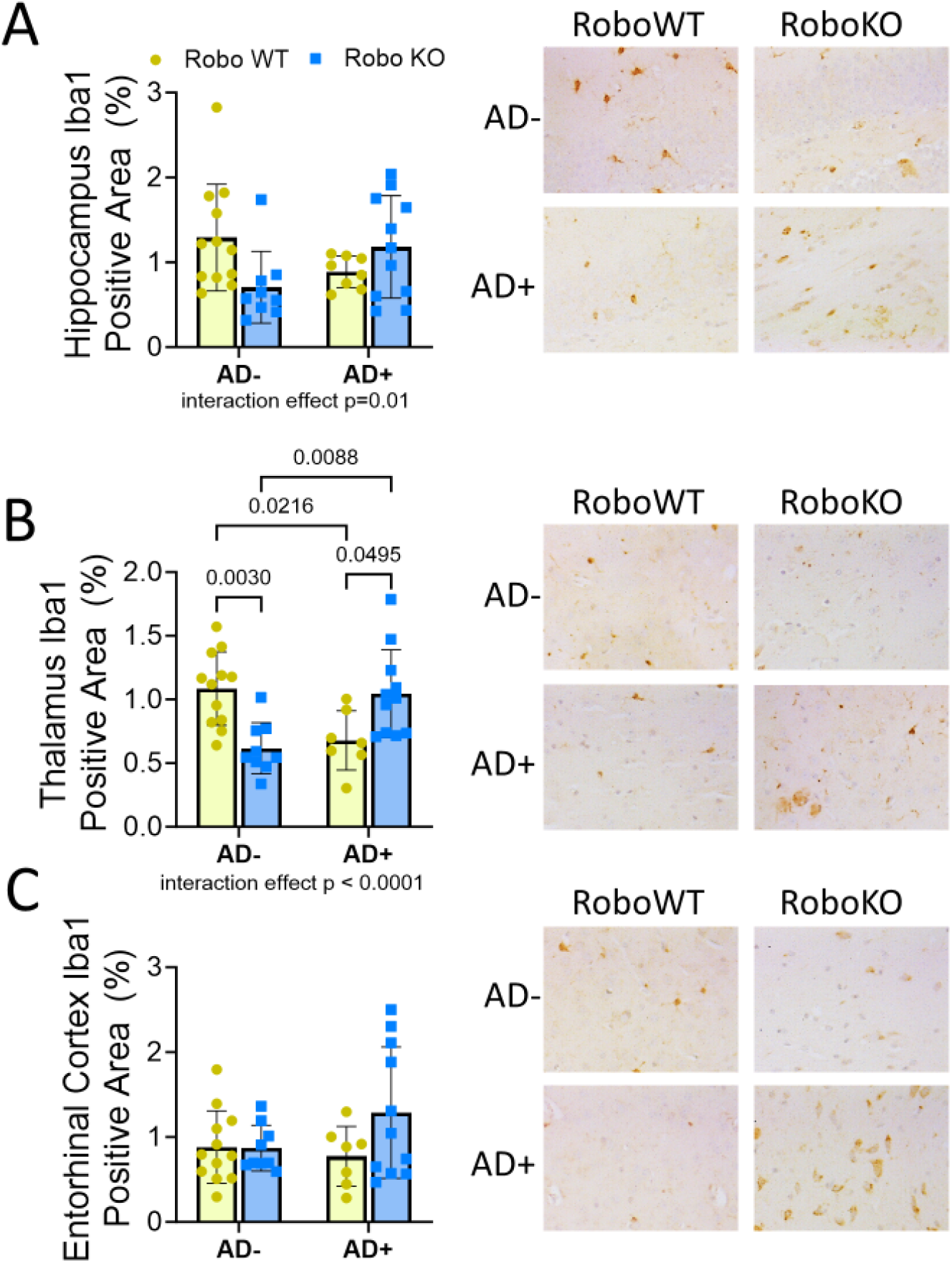
Microglia activation is influenced by both *Robo4* and AD genes. Iba1 quantified by immunohistochemistry in the (A) hippocampus, (B) thalamus, and (C) entorhinal cortex for Robo4 wildtype (WT) or knockout (KO) mice that had mutant Alzheimer’s disease (AD) genes present (+) or absent (-). There was an interaction effect in the hippocampus and thalamus for Iba1 expression (p=0.02 and 0.0004, respectively). There was no effect of *Robo4* or AD genes on Iba1 expression in the entorhinal cortex. Representative images to the right. Values are mean±SD.

### Robo4 deletion impairs motor coordination, but does not affect nest-building or anxiety

Cognitive function, measured by the nest building test, was significantly impacted by AD genotype (main effect: p=0.04), where the presence of AD mutations led to a significant reduction in nest quality (**Figure 3A**). There was no impact of *Robo4* genotype on nest building (p=0.7). Anxiety, measured by the open field test, was not influenced by *Robo4* or AD genotypes (p>0.05) (**Figure 3B**).

**Figure 3.**
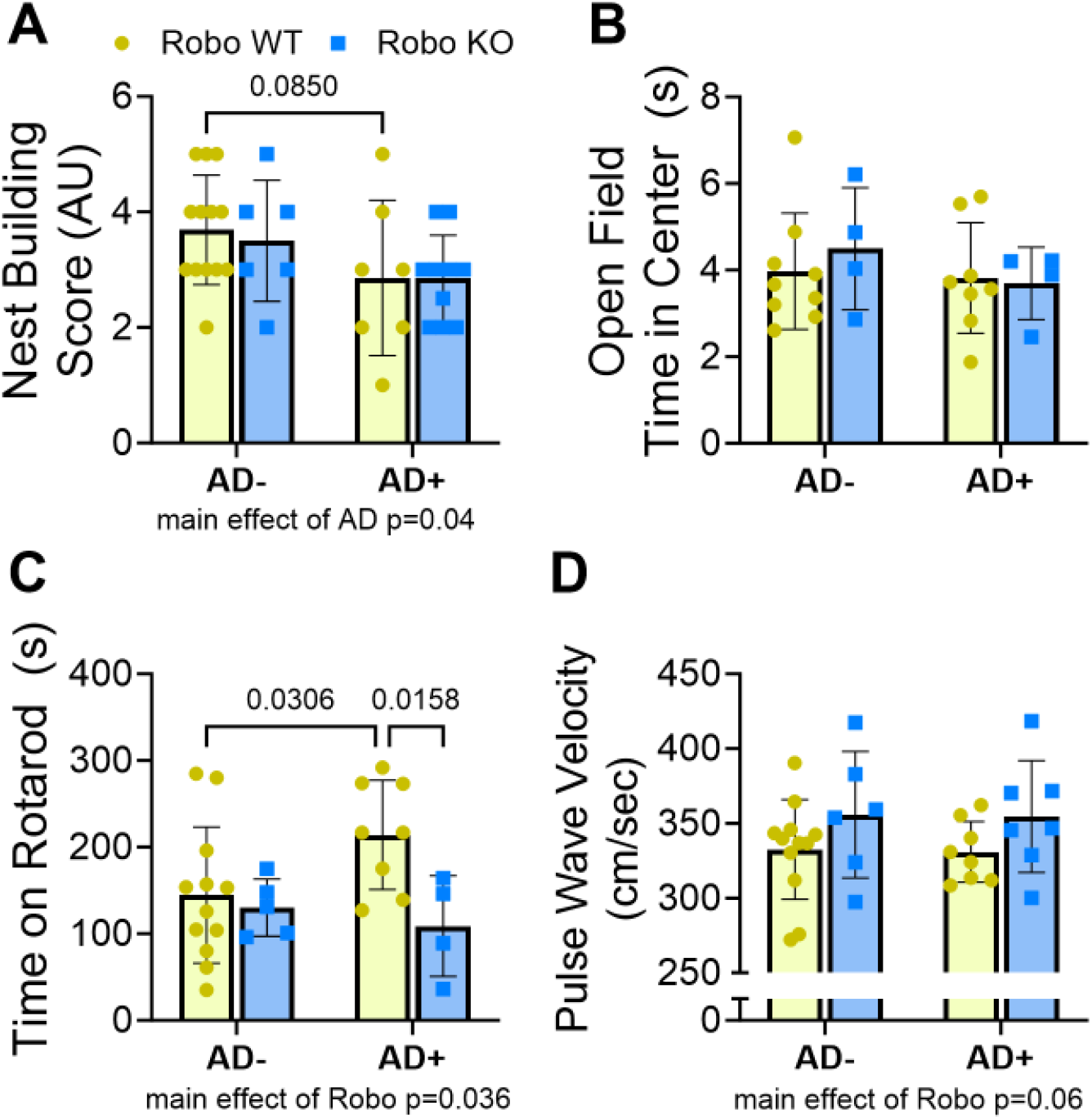
*Robo4* and AD genes influence cognitive and cardiovascular function. We assessed instinctual behavior by nesting building score (A), anxiety during an open field test (B), motor coordination by accelerating rotarod test (C), and aortic stiffness by pulse wave velocity (D) in Robo4 wildtype (WT) or knockout (KO) mice that had mutant Alzheimer’s disease (AD) genes present (+) or absent (-). Instinctual behavior was influenced by AD genes, with AD+ mice having a worse nest building score (p=0.04). There was a main effect of *Robo4* KO on motor coordination (p=0.036) and *Robo4* KO x AD had significantly lower time on the rotarod than *Robo4* WT x AD mice. *Robo4* and AD genes did not influence anxiety or aortic stiffness, however there was a trend for *Robo4* KO mice to have a higher aortic pulse wave velocity (p=0.06). Representative images to the right. Values are mean±SD.

For the accelerating rotarod test, there was a main effect of the *Robo4* genotype (p=0.036), but no effect of AD genotype or an interaction effect (both p>0.05) (**Figure 3C**). The main effect of *Robo4* indicates that the *Robo4* KO animals have worse motor coordination than the *Robo4* WT animals. Further analysis indicated Anecdotally, we also observed atypical behaviors in mice, including continuous jumping and flipping. These behaviors made it impossible to perform other measures of cognitive function, such as a Morris water maze test.

### Trend for greater aortic stiffness with Robo4 deletion

There was no effect of AD genotype on aortic PWV (p=0.9); however, there was a trend for a main effect of the *Robo4* genotype (p=0.06), such that *Robo4* KO mice trend to have higher aortic PWV (**Figure 3D**).

## Discussion

In this study, we demonstrated the impact of Robo4, in combination with AD-related mutations, on neuroinflammation, cognitive function, and arterial stiffness. We found that the knockout of Robo4 led to greater astrocyte activation, as demonstrated by GFAP content. However, the GFAP results did not align with vimentin, another astrocyte marker, which demonstrated an increase in activated astrocytes due to AD mutations, but not Robo4 knockout. We also found that the knockout of Robo4 led to greater activated microglia, as assessed by Iba1, but only in the presence of AD-related mutations. We found that AD mutations, but not Robo4, were associated with cognitive dysfunction by nest building. In contrast, Robo4 deletion, but not AD mutations, was associated with impaired motor coordination. Lastly, Robo4 deletion was associated with greater arterial stiffness, but this trend did not reach statistical significance. In summary, these results demonstrate that Robo4 impacts neuroinflammation, motor coordination, and arterial stiffness, however, the impact on neuroinflammation is dependent on the presence/absence of AD-related mutations and the brain region examined.

### Robo4, neuroinflammation, and BBB permeability

Robo4 prevents neuroinflammation, as demonstrated by our study and others. In our study, we find that Robo4 deletion leads to greater reactive astrocytes and microglia, at least in the thalamus and cortex. These results are in line with previous studies that found that overexpression of SLIT2, the primary Robo4 ligand, leads to fewer reactive astrocytes[11,12]. Previous studies have found that Slit2 impacts neuroinflammation in the hippocampus and cerebral cortex[12], while our study did not find impacts on neuroinflammation in the hippocampus. A potential explanation for the lack of an effect in the hippocampus in our study could be a limited power to detect differences or the broad age range in our study that led to heightened neuroinflammation in even the wildtype mice. In our study, it appears that the presence of AD mutations amplifies the impact of Robo4 deletion on neuroinflammation. Similar to our findings, others have found that the benefits of Robo/Slit are amplified in disease conditions. For example, it was previously found that Slit2 overexpression does not impact microglia activation in middle-aged control (sham) mice[11]. However, in mice who were given cerebral microinfarcts, Slit2 overexpression led to less microglia activation compared with mice without Slit2 overexpression [11]. Thus, it appears that decreases in Slit/Robo signaling lead to neuroinflammation, but it appears that for a substantial impact other stressors need to be present. This suggests that perturbations in Slit/Robo signaling may have minimal impact in young, healthy conditions but may increase their impact in the presence of other stressors.

The mechanism for the impact of Robo4 on neuroinflammation is likely due to the modulation of BBB permeability. A leaky BBB has the potential to allow pro-inflammatory molecules and cells in the blood to enter the brain, thus triggering neuroinflammation[1]. Robo4 is known to control endothelial cell interactions, impacting permeability and sprouting, in established blood vessels[7]. In this way, Robo4/Slit2 signaling leads to increased stability of cerebral endothelial cells and a less permeable blood-brain barrier. For example, after a surgical brain injury in rats, administration of recombinant Slit2 led to less blood-brain barrier permeability compared with untreated rats, and the effects of Slit2 were shown to be mediated by Robo4[13]. The stabilization of the BBB by Robo4 is most likely mediated by an inhibition of VEGF signaling and reducing the pro-permeability effects of inflammatory factors[7]. For example, IL-1beta, TNF, and LPS induce permeability of endothelial barriers, but this is inhibited by Slit2 in a Robo4-dependent manner[8]. However, in the absence of VEGF activation, Robo/Slit2 modulation has minimal effects[7]. As such, it appears that the most striking effects of Robo4 on endothelial barrier permeability will be in the presence of a stressor. Similarly, it was previously shown that Robo4 deletion induces endothelial cell senescence, but only when an additional stressor (in this case iron overload) was present [9]. Thus, the absence of Robo4 can lead to endothelial barrier dysfunction and potentially lead to greater BBB permeability, and this is mostly likely to occur in the presence of an insult that induces pro-angiogenic stimuli, such as aging or brain/vascular injury.

It should be mentioned that the pro-stability effects of Robo/Slit signaling are not entirely consistent in the literature. One previous study found that Slit2 overexpression in wildtype mice leads to greater BBB permeability [14]; however, this study was performed in otherwise healthy mice, and thus, the finding may be due to the lack of a pro-permeability stimulus. In addition, the activation of other Slits or Robos can lead to vascular instability; for example, Slit binding to Robo1 promotes endothelial cell motility, an opposite effect of Robo4 [15]. Further, Slit3 can activate Robo4, and this causes increased angiogenesis, an opposite effect of Slit2 [16]. Similarly, Slit2 can be both pro- and anti-angiogenic, with the pro-angiogenic effects being mediated by mTOR, while the anti-angiogenic effects require ephrin-A1 [17]. We also previously demonstrated that Robo4 has anti-vasodilatory effects, specifically impaired endothelium-dependent vasodilation[18]. This impaired vasodilation may be due to the inhibition of nitric oxide production from VEGF pathways with Robo4 activation. These effects could potentially lead to reduced cerebral blood flow or neurovascular coupling. Thus, while Robo4 appears important for maintaining the BBB and preventing neuroinflammation during pro-angiogenic conditions, this may be context-specific and the impact of Robo4 on vasodilation should be considered.

### VEGF and AD

The impact of VEGF signaling in AD is controversial. Low circulating concentrations of VEGF are associated with AD in clinical studies (https://pubmed.ncbi.nlm.nih.gov/17587256/). Similarly, higher cerebral spinal fluid VEGF is associated with better cognitive function [4]. However, studies of VEGF treatment in mice are inconsistent. One study found that blocking VEGF signaling in AD mice leads to better cognitive function [19], while another study found that VEGF treatment led to better cognitive function in mice [5]. In the context of Robo4, our data and previous studies suggest that the impact of VEGF treatment, whether positive or negative in AD, might depend on the status of Robo/Slit signaling. When proper Robo4 is present to counteract the pro-permeability effects of VEGF, then VEGF may have a beneficial effect [7]. However, in conditions such as aging in females and diabetes [9,10], when Robo4 is known to decline, the impact of VEGF may change.

### Robo4, cognitive function, and motor coordination

The effects of Robo4 signaling on neurocognitive function are inconsistent in the literature. A novel finding from our study is that Robo4 deletion led to impaired motor coordination during an accelerating rotarod test. While the effects of Robo4 on motor coordination have not been previously studied, our results do align with a previous finding demonstrating that Slit3 deletion led to worse rotarod performance [20]. Further studies are needed to determine if the effect of rotarod was neurologic or musculoskeletal.

In our study, we did not find that Robo4 deletion impacted nest building, while the AD-related mutations led to the expected worse performance. In contrast, previous studies have found that Slit2 overexpression, which will activate Robo4, leads to a better performance on the Morris Water Maze, in middle-aged mice and after brain injury [11,12]. The difference between our study and this previous study could be that the Morris Water Maze test is more sensitive to changes in hippocampal dependent memory than nest building. Adding further inconsistencies, another study found that Slit2 overexpression worsens cognitive function, but these studies are in otherwise healthy mice [14]. Our findings that AD-related mutations are associated with worse performance on the nest-building test are consistent with the literature. Thus, the effects of Robo4 on memory are inconsistent in the literature and need further investigation.

### Robo4 and arterial stiffness

We find that Robo4 deletion was associated with a higher aortic stiffness, but this trend did not reach statistical significance. While the effects of Robo4 on arterial stiffness have not been previously studied, it is known that Robo4 impacts the extracellular matrix. For example, Robo4 knockdown increases MMP-9 activity [21]. However, seemingly contradictory to our finding, the knockdown of Slit2 leads to decreased collagen expression in cardiac tissue through Robo1[22]. The tissue of study (vascular vs. cardiac) and Robo isoform may be reasons for the differences between our study and previous findings. Therefore, further studies are needed to understand the mechanisms by which Robo4 may affect stiffness in arteries.

### Limitations

There are a few limitations to this study that should be considered. First, the Robo4 deletion was not specific to cerebral endothelial cells. Therefore, the deletion of Robo4 from endothelial cells outside the brain could impact the outcomes, such as by systemic actions. Second, the Robo4 deletion was constitutive and thus was present during development. However, previous studies have shown that there are no differences in vascular structure observed during development in these mice [7](Jones, Nat Med 2008). Lastly, due to a limited number of mice, we were unable to perform functional BBB permeability studies, therefore we cannot definitively say if the effects we see are due to increased endothelial cell permeability.

### Conclusions

The primary findings from this study are that Robo4 deletion leads to greater neuroinflammation and impaired motor coordination. The presence of AD-related mutations further strengthened the impact of Robo4 deletion, particularly for microglia activation. These results suggest that Robo4 is important for squelching neuroinflammation and maintaining motor coordination in the face of AD. Our findings align with previous studies that show a benefit of Robo4 in maintaining a healthy cerebral vasculature, particularly in the face of injury and/or inflammation. These studies add further evidence in support of targeting the Slit2/Robo4 pathway to limit the damage during AD. However, there remain many unanswered questions about the role of Robo4 in this disease and future studies are required.

## Methods

### Animals

To study the effects of Robo4 in the context of AD, we generated a cross between AD mice [23] with *Robo4* knock-out mice [7] to generate four different genotypes: *Robo4* WT, *Robo4* KO, *Robo4* WT x AD, and *Robo4* KO x AD. *Robo4* KO mice were obtained from Dr. Dean Li’s laboratory at the University of Utah. The AD mice had a knock-in of the amyloid precursor protein Swedish mutation and the tauP301L mutation, obtained from the Jackson Laboratory. Male and female mice were studied at young (6 mo), middle-age (12-14 mo), or old (22-24 mo) ages. We found no age or sex differences for any outcome within each genotype, and thus, we combined the age and sex groups for each genotype. After collecting behavior measurements, mice were euthanized by cardiac puncture and perfusion with saline. Following saline, mice were perfused with 4% paraformaldehyde to preserve cerebral tissue. All mice were housed in an animal care facility on a 12/12 h light-dark cycle at 24°C. All animal procedures conformed to the *Guide to the Care and Use of Laboratory Animals* (8^th^ version) and were approved by the Institutional Animal Care and Use Committee at the University of Oregon.

### Aortic Stiffness

We measured aortic stiffness using pulse wave velocity. Mice were anesthetized using 3% isoflurane in 100% oxygen and then placed in the supine position on a temperature-controlled surgical pad at 37°C (Indus Instruments, Webster, TX, USA). ECG was recorded throughout the procedure and two Doppler Transducers were positioned at the aortic arch and abdominal aorta to collect waveforms simultaneously. After recording 3 ∼10 s intervals of the Doppler signal, the distance between the two probes was measured. To analyze the difference in pulse arrival times the Doppler Signal Processing Workstation program (Indus Industries, Webster, TX, USA) was used. Two blinded researchers analyzed each file independently, and inter-observation confidence intervals were measured; those over 15% were reanalyzed by a third researcher. PWV was calculated dividing the distance (in cm) by ejection time recorded between the two Doppler Transducers.

### Cognitive Function

Nest Building: To test for changes in instinctual behavior, we conducted a nest-building test. Mice were singly housed during the dark cycle for 12 h in a cage containing a cotton nestlet and no other enrichment. At the end of the 12 h, the cotton nestlet was scored against a preset scale from 0-5, with 0 indicating an untouched nestlet and 5 indicating a fully built nest [24].

Open Field: To test for changes in anxiety levels we conducted the open field test, as previously described [25]. Briefly, mice were placed in an enclosed, empty arena (40cm by 40 cm) and allowed to explore for 10 minutes, during this time a video was recorded using Ethovision software (version 12, Noldus, Leesburg, VA, USA). Analysis of open field was conducted with Ethovision and exploration criteria were defined as the amount of time the mouse spent in a predetermined inner square (20 cm by 20 cm) of the area versus the outer perimeter of the arena, where mice that spent more time in the inner square were considered to have less anxiety.

Rotarod: To test for changes in motor coordination we conducted the Accelerating Rotarod test. On day 1 of the test, mice were placed on the Rotarod and remained on the Rotarod at 4 rpm for at least 90 s to acclimate. On day 2 of the test, mice performed 3 trials where the rotation speed increased from 4 rpm to 40 rpm over 300 seconds. The time for the trial was recorded as when the mouse fell off the Rotarod or hung on for two rotations or when 300 s passed. The animals received a 10-minute rest between each trial. The average of the three trials is reported [26].

### Histology

After paraformaldehyde perfusion, brains were incubated in 4% paraformaldehyde for 48 h and then transferred to distilled water for longer-term storage. Brains were embedded in paraffin before being cut at 20 um thickness. Immunohistochemical staining was conducted as described previously [27], using primary antibodies for Iba1 (1:5000, ProteinTech, 10904-1-AP), GFAP (1:500, ProteinTech, 16825-1-AP), and vimentin (1:500, ProteinTech, 10366-1-AP), a rabbit secondary antibody, and DAB. Analysis of histology was conducted by first imaging the regions of interest (hippocampus, entorhinal cortex, and thalamus) using a Leica DMi1 microscope. Image J software (NIH, Bethesda, MD, USA) was used for analysis of percent positive area.

### Statistical Analysis

Statistical analyses were performed with GraphPad Prism 10.0. A one-way analysis of variance (ANOVA) between different ages (young, middle, and old age) and between sexes within each genotype was conducted initially to determine if groups could be combined. There were no significant differences found, and thus, the age groups were combined. Outliers were determined to be values with Z-scores outside the range of -2.0 to 2.0 and were removed from the data set. All groups were normally distributed as assessed by a Shapiro-Wilk test. If normality was not met, data was transformed by taking the square root of each sample. The Shapiro-Wilk test was used again to confirm normality was met if the data was transformed. A two-way ANOVA (AD genotype x Robo4 genotype) was conducted for all outcomes. In cases of a significant F value, post hoc analyses were performed using the Tukey correction for preplanned comparisons. Significance was set at p < 0.05 and all values shown are means ± standard deviation.

