## Supplemental Data for "Deletion of Robo4 worsens neuroinflammation and motor coordination in a mouse model of Alzheimer’s disease"

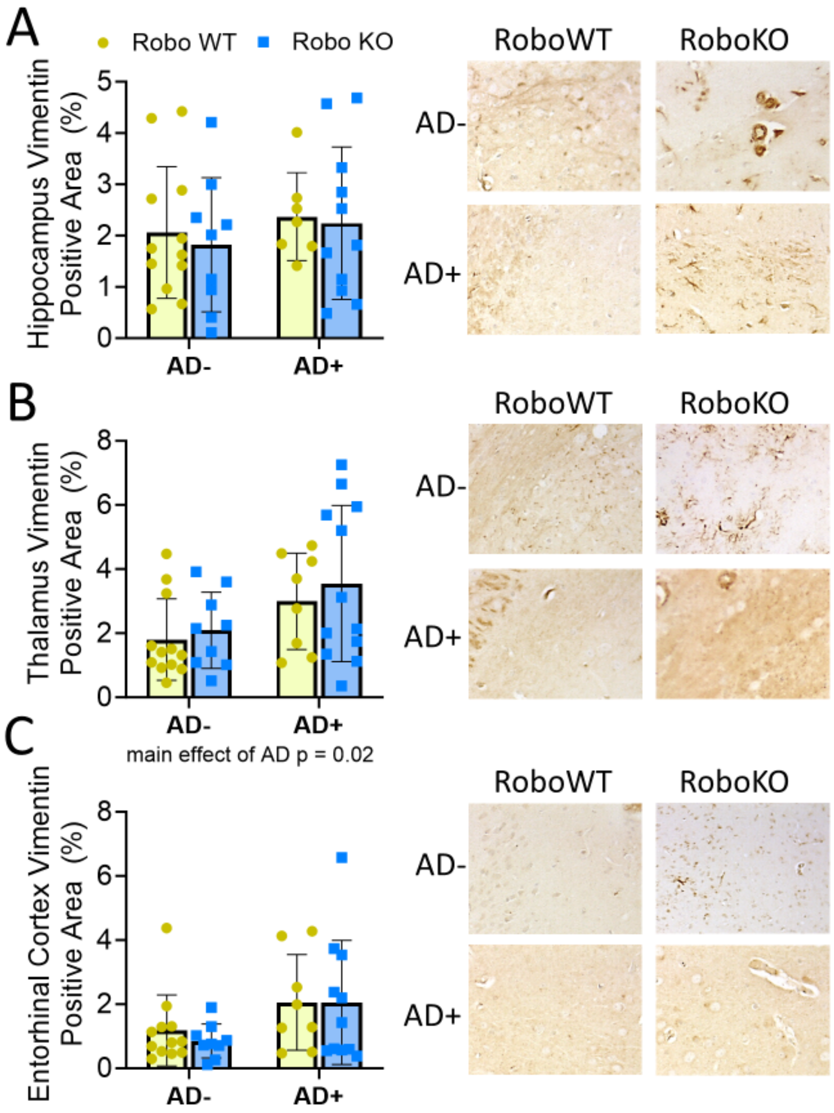


**Supplemental Figure 1. AD genes influence vimentin expression in the thalamus.** We assessed vimentin expression in the hippocampus (A), thalamus (B), and entorhinal cortex (C). There was a main effect of AD genes in the thalamus. Vimentin expression data from the hippocampus and entorhinal cortex did not pass normality and was not analyzed, data are shown for visual representation. Representative images to the right. Values are mean±SD.
